# Molecular signatures of sexual communication in the phlebotomine sand flies

**DOI:** 10.1101/2020.08.11.247155

**Authors:** Paul V. Hickner, Nataliya Timoshevskaya, RJ Nowling, Frédéric Labbé, Andrew D. Nguyen, Mary Ann McDowell, Carolina N. Spiegel, Zainulabeuddin Syed

## Abstract

Phlebotomine sand flies employ an elaborate system of pheromone communication wherein males produce pheromones that attracts other males to leks (thus acting as aggregation pheromone) and females to the lekking males (sex pheromone). In addition, the type of pheromone produced varies among populations. Despite the numerous studies on sand fly chemical communication, little is known of their chemosensory genome. Chemoreceptors interact with chemicals in an organism’s environment to elicit essential behaviors such as the identification of suitable mates and food sources, thus, they play important roles during adaptation and speciation. Major chemoreceptor gene families, odorant receptors (ORs), gustatory receptors (GRs) and ionotropic receptors (IRs) together detect and discriminate the chemical landscape. Here, we annotated the chemoreceptor repertoire in the genomes of *Lutzomyia longipalpis* and *Phlebotomus papatasi*, major phlebotomine vectors in the New World and Old World, respectively. Comparison with other sequenced Diptera revealed a large and unique expansion where over 80% of the ~140 ORs belong to a single, taxonomically restricted clade. We next conducted comprehensive analysis of the chemoreceptors in 63 *Lu. longipalpis* individuals from four different locations in Brazil representing allopatric and sympatric populations and three aggregation-sex pheromone types (chemotypes). Population structure based on single nucleotide polymorphisms (SNPs) and gene copy number in the chemoreceptors corresponded with their putative chemotypes, and corroborate previous studies that identified multiple populations. Our work provides genomic insights into the underlying behavioral evolution of sexual communication in the *L. longipalpis* species complex in Brazil, and highlights the importance of accounting for the ongoing speciation in central and South American *Lutzomyia* that could have important implications for vectorial capacity.

## Introduction

Globally, vector-borne diseases account for more than 17% of all infectious diseases every year. One such disease, leishmaniasis, is endemic in 98 countries with an estimated 700,000 to 1 million new cases leading to 26-65,000 deaths each year[1]. Leishmaniasis is a group of vector-borne diseases caused by protozoan parasites from the genus *Leishmania* and is considered among the most important neglected tropical diseases [2]. Major vectors of this diseases are *Phlebotomus* spp. in the Old World and *Lutzomyia* spp. in the New World. *L. longipalpis* is found in a wide but discontinuous geographical distribution from Mexico to Argentina, where they inhabit diverse ecological environments, while *Phlebotomus* has a wide geographical distribution (from southern Europe, northern Africa, the Middle East, and India) and inhabits a variety of ecological niches from tropical climates to arid desert [3].

Sand flies of both the genera display robust olfactory behaviors to locate suitable hosts, oviposition sites and mates [4]. In contrast to most disease vectors, which do not employ long range chemical communication to locate potential mates [5], sand flies of the *Lutzomyia* species complex employ an elaborate pheromone communication system [4, 6], wherein males produce pheromone(s) that attract conspecific males to courtship aggregations (leks), and attract females to the lekking males. In *L. longipalpis*, these aggregation-sex pheromones are produced in tergal glands that appear as pale patches or “spots” on the abdomen [6]. Historically, the number of spots — one spot (1S) or two spots (2S) — served as a potential phenotypic marker for cryptic species complex in *L. longipalpis* [6]. The first evidence of the existence of the *L. longipalpis* species complex was obtained in Brazil [7], and genetic variability in sand flies with potential implication in leishmaniasis has long been emphasized [8]. Different aggregation-sex pheromones have been described from male *L. longipalpis* from Brazil: *S*-9-methylgermacrene-B (9MGB), (1*S*,3*S*,7*R*) 3-methyl-α-himachalene (3MαH), two cembrene isomers (Cemb-1 and Cemb-2), and a novel chemotype based on variation in the quantity of 9MGB produced (9MGB+) [9, 10]. Cemb-1 was recently reclassified as a novel diterpene and named sobralene [11]. In addition to pheromone communication, *L. longipalpis* males produce a song during copulation—an acoustic signal generated by vibrating their wings—that varies among populations and that can be broadly categorized as “pulse-type” or “burst-type”. Variation in copulation song and aggregation-sex pheromone, together with subsequent crossing studies and genetic differentiation, have provided compelling evidence for *L. longipalpis* comprising a species complex [6, 9].

Signaling and reception evolve in synchrony and shape the resulting behavior. Broadly defined as ‘sensory drive’, especially in the context of environmental conditions [12], this process is prominent in chemosensation across phyla [13]. Genomic changes underlie behavioral evolution; thus, studies of sensory systems and their genetic correlates provide insights into the patterns of ecological evolution [14]. This is particularly evident in arthropod vectors that transmit various life-threatening diseases [15–17]. For example, multiple anopheline species altered their behavioral repertoire following sustained use of bed nets and indoor spraying[18]; ancestral African aedenine populations evolved to be human commensals facilitated by behavioral and genetic changes [19, 20] such as house-entering behavior [21], and enhanced human preference [19] facilitated by chemoreceptor gene families [19, 22, 23]. Since the communication system – comprising songs and pheromones -- is well defined in sand fly species complex, we undertook a study to isolate and identify the genetic basis of such behaviors.

Here we report a molecular evolutionary analysis of the sand fly chemoreceptor genomes comprising the ORs, GRs, and IRs, which are among the largest gene families, and together define the reception and perception of odors associated with hosts, mates and oviposition sites in insects [24, 25]. We annotated the chemoreceptors in the whole genome assemblies of two phlebotomine sand flies, *L. longipalpis* and *P. papatasi*, and conducted comparative analyses with other Diptera. In addition, we investigated variation in the chemoreceptor genomes of 63 *L. longipalpis* individuals collected from four locations in Brazil, representing sympatric and allopatric populations and three different aggregation-sex pheromones. Investigations into the molecular signatures of genetic variation have traditionally focused on single nucleotide polymorphisms (SNPs) [26], but recently there has been an emphasis on gene copy number variation (CNV) as an additional source of genomic variation in the insect chemoreceptors [25, 27]. Our analysis of SNPs and CNV provides novel insights into the evolution of chemosensation, thus providing a framework for future studies on the molecular basis of chemical communication in the phlebotomine sand flies.

## Methods

### Annotation of the chemoreceptor repertoires

Genes were manually annotated as described previously [28]. Briefly, genomic loci encoding odorant receptors (ORs), gustatory receptors (GRs) and ionotropic receptors (IRs) were identified by tBLASTn analysis of the *L. longipalpis* (LlonJ1) and *P. papatasi* (PpapI1) genome assemblies in VectorBase [29] using *Anopheles gambiae* and *Drosophila melanogaster* peptides as queries. All BLAST analysis was conducted using the BLOSM62 scoring matrix and a maximum E value of 10000, with a tBLASTn word size of 3 and a BLASTn word size of 11. *An. gambiae* gene models were downloaded from VectorBase (AgamP4.9), and *D. melanogaster* gene models were downloaded from FlyBase (release FB2017_05). An exhaustive screen of the *L. longipalpis* and *P. papatasi* genome scaffolds was performed by reciprocal BLAST analysis using the sand fly chemoreceptor gene models. Genes were prefixed with either Llon (*L. longipalpis*) or Ppap (*P. papatasi*). The ORs and GRs were numbered arbitrarily with the following exceptions: the putative CO2 receptors were numbered Gr1 and Gr2, followed by the sugar receptors; and several, but not all, 1:1 orthologs between *L. longipalpis* and *P. papatasi* were numbered the same (e.g. Gr1, Gr2, Gr13, Gr26). Most IRs were named based on homology to *D. melanogaster*, while Ir101 and Ir102 were named based on their homology to Ir101 in *An. gambiae*.

Genes with gaps, premature stop codons and indels that were “fixed” (when possible) by BLASTn analysis of the SRAs were suffixed with “_fix”. Indels and premature stop codons that could not be fixed or were confirmed with SRAs were considered pseudogenes and suffixed with “P”. Genes spanning two or more different scaffolds were suffixed with “_join”, and genes that were on the same scaffold but on different strands were suffixed with “_strand”. Genes annotated from the *de novo* assemblies are suffixed with “_denovo”. Partial gene models encoding ≤330 amino acids were omitted from the dataset.

### Phylogenetic analysis

A species phylogeny was estimated to illustrate the evolutionary relationships among the Diptera used for comparative analysis of chemoreceptor repertoire size. The OrthoFinder v2.3.3 program was used for orthologous group selection among peptides in *Anopheles gambiae, Aedes aegypti, Culex quinquefasciatus, P. papatasi*, *L. longipalpis*, *Mayetiola destructor*, *Musca domestica*, *D. melanogaster, Glossina morsitans* and the outgroup *Bombyx mori* [30]. The species tree was estimated with OrthoFinder using the FastME distance-based program [31].

OR, GR and IR gene trees were estimated by first conducting multiple sequence alignments of *An. gambiae, M. destructor, P. papatasi, L. longipalpis* and *D. melanogaster* peptides [32–36] using Muscle v3.8.31 [37]. The multiple sequence alignments were trimmed using the *automated1* option in Trimal v1.4 [38]. Maximum likelihood trees were inferred using the JTT model of protein substitution in RAxML v.8.2.4, which was chosen based on *protgammaauto* model selection [39]. Branch support was estimated using 500 bootstrap replications. The OR, GR and IR trees were rooted with *Orco*, the CO_2_ receptor clade and the *Ir8a/Ir25a* clade, respectively.

### Analysis of *L. longipalpis* populations in Brazil

The SRA toolkit v2.9.2 [40] was used to download fastq files from NCBI containing paired-end reads (101 bp x 2) for 63 *L. longipalpis* individuals from Jacobina (n = 14), Lapinha Cave (n = 11), Marajó (n = 9) and Sobral (n = 29) Brazil. Accession numbers for SRA downloads are reported in Supplemental Table S1. The individuals from Sobral were further grouped based on the number of spots on the abdomen — Sobral with one spot (1S, n = 13) and Sobral with two spots (2S, n = 16). Due to the number of fixes made to the gene models, several redundant genes, extraneous fragments and six missing genes, we used BEDtools v2.28.0 [41] to hard-mask all chemoreceptor loci in the LlonJ1 genome assembly, then appended the manually curated OR, GR and IR gene regions. This “revised” assembly (Supplemental data, rev_assembly.fa) was used for subsequent analyses. Chemoreceptor gene regions comprised introns, exons and 300-500 bp flanking regions (when possible). We used BWA mem v0.7.17 [42] to map the reads to the revised genome assembly. PCR duplicates were removed using SAMtools v1.9 [43]. Reads with soft-clipping on both ends (marked with S in CIGAR string) were removed using custom python scripts. Single nucleotide polymorphisms (SNPs) were called using BCFtools [43]. Sites with a PHRED score <30 and sequencing depth below 0.5 and above 1.5 modal coverage for that individual were omitted. For example, sites below 25X and above 75X were omitted in an individual with a 50X modal coverage.

Phylogenetic relationships among the field isolates from Brazil were estimated with 18,254 SNPs in the exons of all 100 single-copy orthologs using the neighbor joining method in TASSEL v5.2.57 [44]. The principal component analysis (PCA) approach in PCadapt [45] was used to identify chemoreceptors associated with differences in chemotype and/or population structure. We limited the analysis to SNPs (n=18,254) in the exons of genes that were single-copy in all 63 individuals (n=100). Preliminary analysis with *K*=20 and a minimum minor allele frequency of 0.1 was used to select the appropriate number of *K* (principal components) for subsequent analyses, which was determined to be *K*=5 based on the scree plot (Fig S4A). Component-wise analysis with *K*=5 and a minimum minor allele frequency of 0.1 resulted in 6,759 SNPs passing criteria and 502 outlier SNPs after Bonferroni correction (qvalue<0.05) for multiple tests (Fig S4B). Due to the large number of genes with only 1 or 2 outlier SNPs, we highlighted the top three genes with the highest number of SNPs based on SNPs per kb of CDS. The entire list of outlier SNPs and their associated PC can be found in.

We used the program ADMIXTURE v1.3 [46] to estimate genetic introgression among populations and chemotypes. ADMIXTURE was run for *K* 1 through 7 (number of ancestral populations) with 5-fold cross-validation. Each ADMIXTURE analysis was repeated 30 times with different seeds, resulting in a total of 210 runs. To better understand the different solutions reported by ADMIXTURE, for each value of *K* we compared solutions and produced a major cluster of solutions that give similar results using the online version of CLUMPAK with default settings [47]. We used the ADMIXTURE cross-validation procedure to estimate the most preferable number of *K* to provide a complementary view to the Bayesian clustering analysis independently of any model assumptions [48].

The Tablet software v1.19.05.28 program [49] was used to visualize sequence read alignments, which revealed a number of apparent gene absences/losses. To estimate the extent of copy number variation in 245 chemoreceptor genes in all 63 individuals (~15,000 genes) we used background normalized sequencing depth to estimate gene copy number (CN). Read depth at each position in the chemoreceptor exons was extracted using SAMtools utility *depth* with option *-a* to take in account bases with zero depth of coverage [43]. The mean read depth for each chemoreceptor gene was normalized to modal depth across all exons as follows:

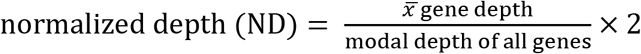

Gene CN was calculated by rounding normalized depth (ND) to the nearest whole number (CN =||ND||). Modal depth ranged from 35 to 133 (mean 67.8) across individuals. A heatmap with hierarchical clustering (one minus the cosine similarity with complete linkage) using CN of chemoreceptor genes in all 63 individuals was calculated using the Morpheus software program (https://software.broadinstitute.org/morpheus).

PCA of normalized sequencing depth was conducted using the covariance method in SigmaPlot v14.0 (Systat Software, San Jose, CA). The one-way analysis of variance and Holm-Šídák test for post-hoc pairwise analysis was conducted using SigmaPlot v14.0 to test for differences in the mean number of absent (CN = 0), single-copy (CN = 2), and duplicated (CN >2) gene lineages among and between chemotypes. Pairwise V_ST_, a measure of population differentiation analogous to F_ST_ but based on CNV, was calculated using methods described previously [50]. For pairwise analyses of V_ST_, populations were combined according to their putative aggregation-sex pheromone (chemotype). Specifically, Marajó, Sobral 2S and Jacobina-B (sobralene); Sobral 1S-A and Lapinha (9MGB); and Sobral 1S-B and Jacobina-A (3MαH).

To evaluate ND as a proxy for CN, we used Megahit [51] to construct a *de novo* genome assembly for each individual with the parameters --min-count 2, -- k-min 21, --k-max 141, --k-step 10. We annotated a subset of the chemoreceptors in the *de novo* assemblies using Geneious v6.1.8 (https://www.geneious.com) for BLAST analysis and methods previously described for the reference genomes. Seven genes (Or67, Or115, Or116, Or109, Or137, Or138 and Or139) were annotated or confirmed missing/nonfunctional in the *de novo* assemblies of all 63 individuals. Genes and alleles that are full length based on read coverage and/or manual annotation are presumed to be functional are referred to as “intact”.

## Results

The sand fly OR repertoires are relatively large and uniquely expanded among the dipterans analyzed here, with 140 in *L. longipalpis* and 142 in *P. papatasi* (Fig 1A). Over 80% of the sand fly ORs belong to a single, taxonomically restricted lineage with no close relationship to other ORs included in our phylogenetic analysis (Figs 1B and S1). Only 5 lineages were conserved throughout the Diptera suggesting that gene death through pseudogenization and/or deletion has helped shape the sand fly repertoire of ORs (Fig S1). One notable difference between the two sand fly species is the *AgamOr1-DmelOr46a* lineage, which is lost in *L. longipalpis* but expanded to five copies in *P. papatasi* (Fig S1). The GR repertoires include 82 genes in *L. longipalpis*, and 77 in *P. papatasi* (Figs 1C and S2). Interestingly, in contrast to most Diptera which have three CO_2_ receptors conferring the detection of this important host odor cue (25), only two intact genes (*Gr1* and *Gr2*) were found in the sand fly genomes. Screening of sequence read archives (SRA) for the third CO_2_ receptor revealed a highly degraded pseudogene in each sand fly species, thus suggesting the loss of *DmelGr63a/MdesGr3/AgamGr24* ortholog in a common ancestor. The 3^rd^ major chemosensory gene family, the IRs, was smaller (23 and 28 IRs in *L. longipalpis* and *P. papatasi*, respectively) compared to mosquitoes and *D. melanogaster* (Figs 1D and S3). Of the three chemoreceptor families analyzed, the IRs were the least dynamic based on the number and extent of expanded and lost lineages. Coding sequences for ORs, GRs and IRs in *L. longipalpis* and *P. papatasi* are provided in Supplemental Table S2.

**Fig. 1.**
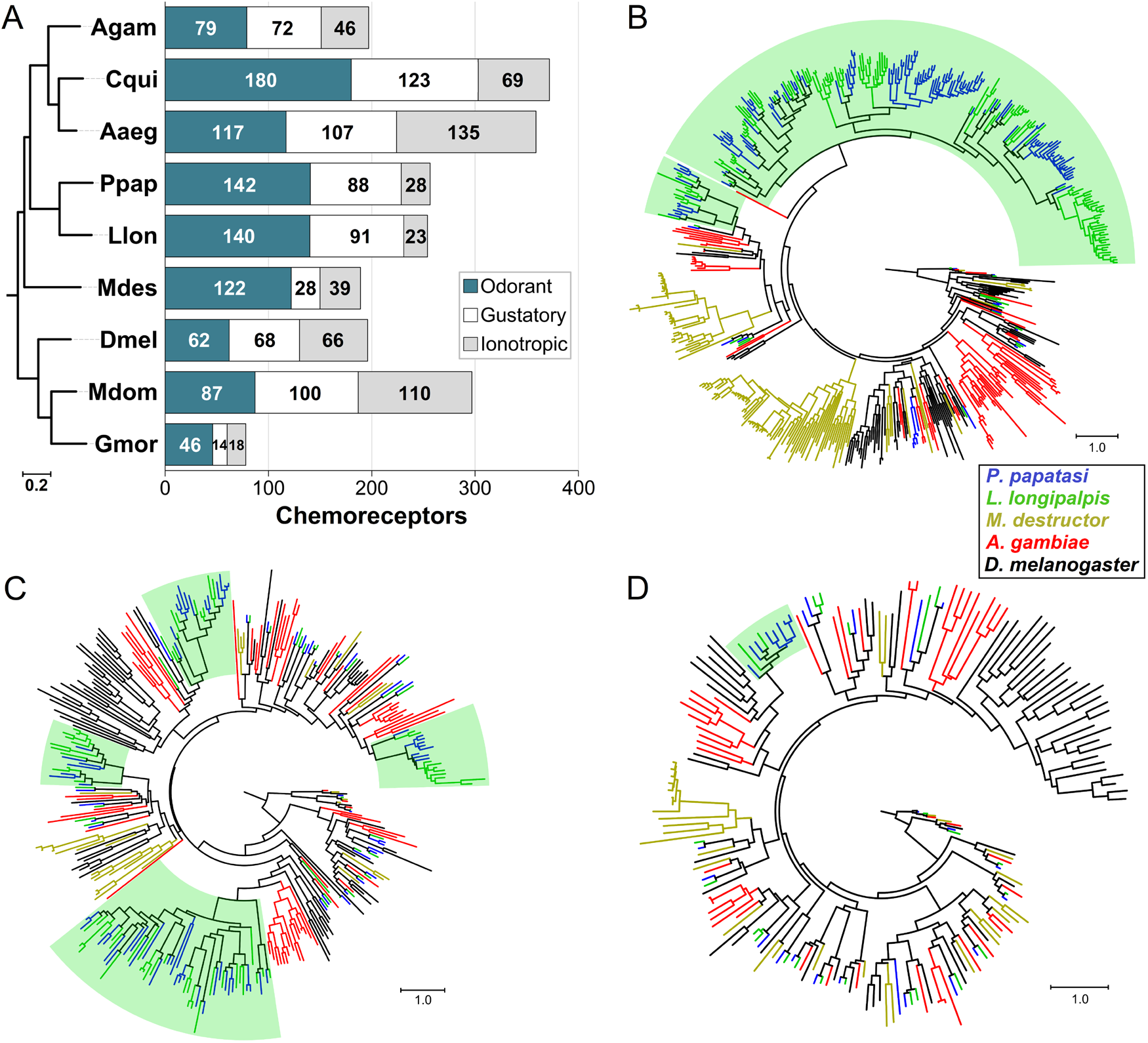
Chemoreceptors in the phlebotomine sand flies. **(A)** Chemoreceptor repertoire size in the sand flies *L. longipalpis* (Llon) and *P. papatasi* (Ppap) compared with *An. gambiae* (Agam), *C. quinquefasciatus* (Cqui), *Ae. aegypti* (Aaeg), *Ma. destructor* (Mdes), *Mu. domestica* (Mdom), *G. morsitans* (Gmor) and *D. melanogaster* (Dmel). Species tree estimated using OrthoFinder with the multiple sequence alignment option and 252 single-copy orthologs. Phylogenetic analysis of chemoreceptors in five Diptera revealed a large taxonomically-restricted clade comprising over 80% of the *L. longipalpis* and *P. papatasi* ORs (highlighted in green) **(B)**. Several smaller lineage expansions are evident in the GRs **(C)**, while only one IR lineage (Ir7c) was expanded in the sand flies **(D)**. Phylogenies were estimated using *L. longipalpis, P. papatasi*, *A. gambiae, M. destructor* and *D. melanogaster* protein sequences aligned with ClustalX. The JTT model of protein substitution and Maximum Likelihood method in RAxML v.8.2.4 were used for tree estimation. The trees are rooted at the branch leading to Orco, the CO_2_ receptors, and the Ir25a and Ir8a clades for ORs, GRs and IRs, respectively. Branch support based on 500 bootstrap replications. Scale bars indicate the number of amino acid substitutions per site.

New World sand flies in the *L. longipalpis* complex employ an exquisite communication system, whereby males produce pheromones and copulatory songs to identify their mates. Phylogenetic analysis based on SNPs (n=18,254) in the single-copy chemoreceptor loci (n=100) in 63 *L. longipalpis* individuals from four sites in Brazil (Fig 2A) revealed distinct clades that grouped individuals and populations broadly on the basis of chemotypes and copulation songs. The most prominent separation was between the ‘burst type’ clade comprising Marajó, Sobral 2S and 6 Jacobina individuals with undetermined song type, and the remaining into ‘pulse type’ composed of Lapinha, Sobral 1S and 8 Jacobina (Fig 2B).

**Fig. 2.**
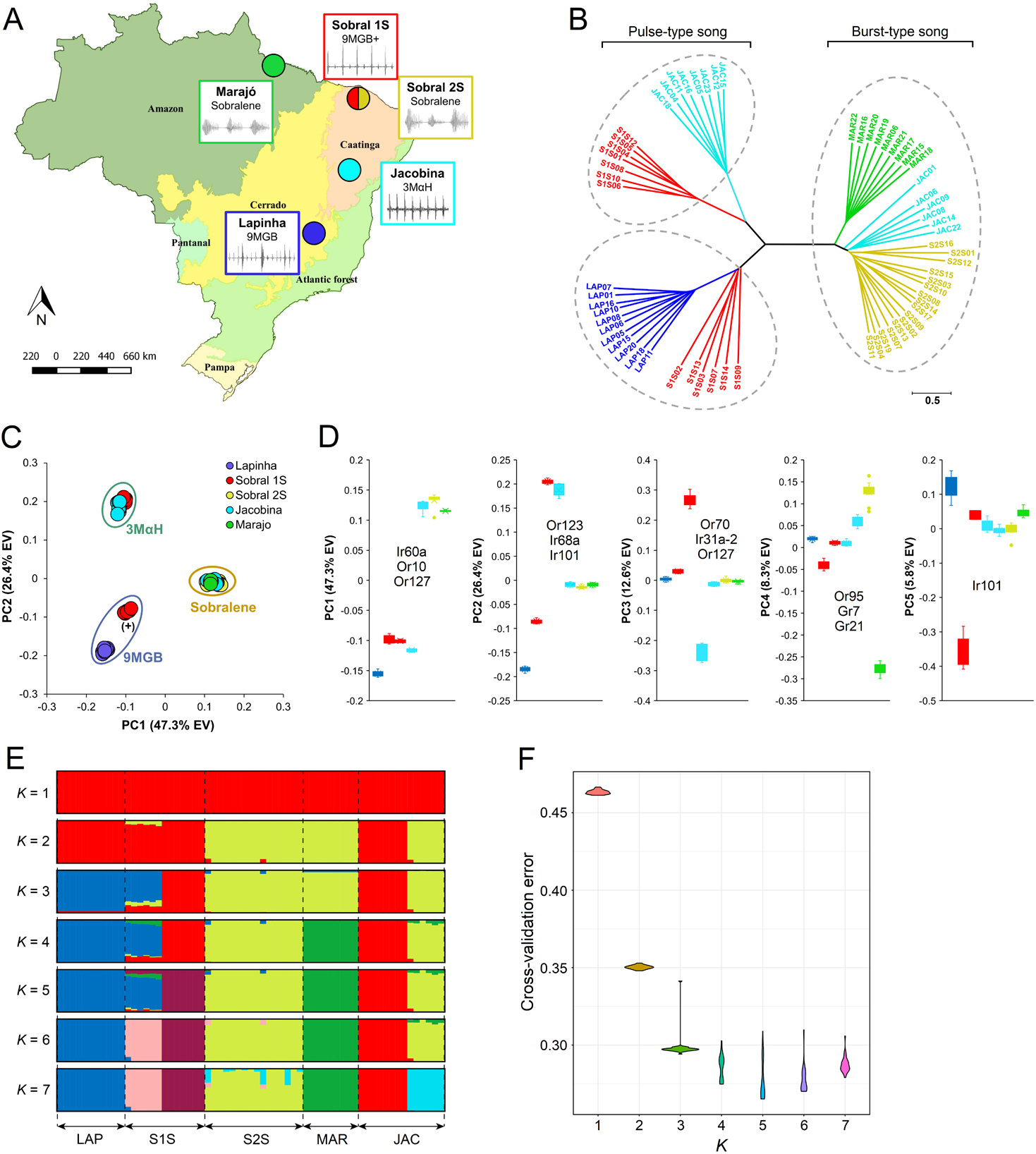
Population structure of 63 *L. longipalpis* from four sites in Brazil based on SNPs in 245 chemoreceptor genes. **(A)** Geographic distribution, aggregation-sex pheromones (9MGB, 3MαH, sobralene) and copulatory songs based on previous studies of *L. longipalpis* in Brazil (16). **(B)** Individuals from Sobral (with 2 spots: S2S), Marajo and Lapinha formed discrete clades, while individuals from Sobral (with one spot, S1S) and Jacobina split into two clades each. Unrooted tree estimated using SNPs in the exons of all 100 single-copy orthologs and neighbor-joining method in Tassel v5.2.57. **(C)** Principal component analysis (PCA) was used to identify loci associated with population structure and conducted using *pcadapt* (explained variance EV). The first two principal components accounted for 73.6% of the total variation and grouped individuals into four clusters. Putative chemotypes were assigned based on previous studies and PCA clustering patterns. **(D)** Principal components 1-5 and genes with the highest number of SNPs based on component-wise outlier analysis in *pcadapt*.

We used PCA to further analyze the population structure and to identify chemoreceptors significantly associated with population differentiation. The first two principal components explained 73.6% of the total variation and grouped individuals into three discrete clusters (Fig 2C) that represented three putative pheromone types (chemotypes): a sobralene cluster composed of all individuals from Marajó and Sobral 2S, and 6 Jacobina (hereafter, JAC-B); a 3MαH cluster comprised of 7 Sobral 1S (hereafter, S1S-B) and 8 Jacobina individuals (hereafter, JAC-A). Interestingly, a third cluster of 9MGB was apparent that included 6 Sobral 1S (hereafter, S1S-A) and all Lapinha individuals. This serendipitous grouping is consistent with the reported findings by Hamilton et al. who classified Lapinha and S1S-A as different chemotypes based on quantitative differences in 9MGB producing individuals [10]. These populations were named 9MGB (Lapinha) or 9MGB+ (S1S-A). With over 47% of the total variation captured, PC1 broadly separates individuals based on the major song types (Fig 2C). It will be exciting to correlate chemotype separation from our analyses with the estimated divergence of burst and song which occurred ca. 0.5 – 0.7 mya [9, 52].

Having found discrete chemotype clusters based on SNPs, we aimed to identify genes contributing to the observed patterns using PCadapt [45]. We identified 502 SNPs, of which 164, 147, 170, 13 and 8 were associated with PC1 through PC5, respectively (Supplemental Table S3). Three genes with significant contribution, defined as ‘genes with greatest number outlier SNPs’ in *pcadapt*, included *Ir60a, Or10* and *Or127* that were involved in the separation of Sobralene from the 9MGB and 3MαH chemotypes (Fig 2D). Or123, Ir68a and Ir101 were involved in the separation between the 9MGB and 3MαH chemotypes in PC2 (Fig 2D). PC3 through PC5 separated populations, where JAC-A and S1S-B (3MαH) were separated by PC3, Marajó was separated from all others by PC4, and S1S-A and Lapinha were separated by PC5. Intriguingly, four of the 11 genes with the highest number of SNPs underscoring PC1-5 were IRs, which are increasingly being implicated in multimodal signaling. Admixture analysis revealed seven groups that were clearly distinguishable at K = 7, consistent with the PCA and ML tree (Fig 2E). However, a clear modeling choice for the number of K was not indicated by cross-validation error analysis, which suggested K is between 3 and 7 (Fig 2F). Despite this limitation, little introgression was indicated, with most being among the sympatric populations in Sobral and Jacobina (Fig 2E).

While SNP analysis revealed population structuring that is largely consistent with previously described chemotypes, we noticed a surprisingly large number of missing genotypes, which prompted us to investigate potential CNV. Variation in gene copy number has been hypothesized to play an important role in the emergence of adaptive traits within and among populations [53]. Analysis of the read coverage revealed extensive variation in depth of coverage across loci (Fig 3A). For estimation of gene copy number (CN), we calculated a normalized sequencing depth based on the modal coverage of the gene coding regions of all 15,435 loci (63 individuals x 245 genes) (Fig 3B and Supplemental Table S4). Normalized depth (ND) ranged from 0 to 6.56 with a distribution having a central tendency ~2 in all three chemotypes (Fig 3C).

**Fig. 3.**
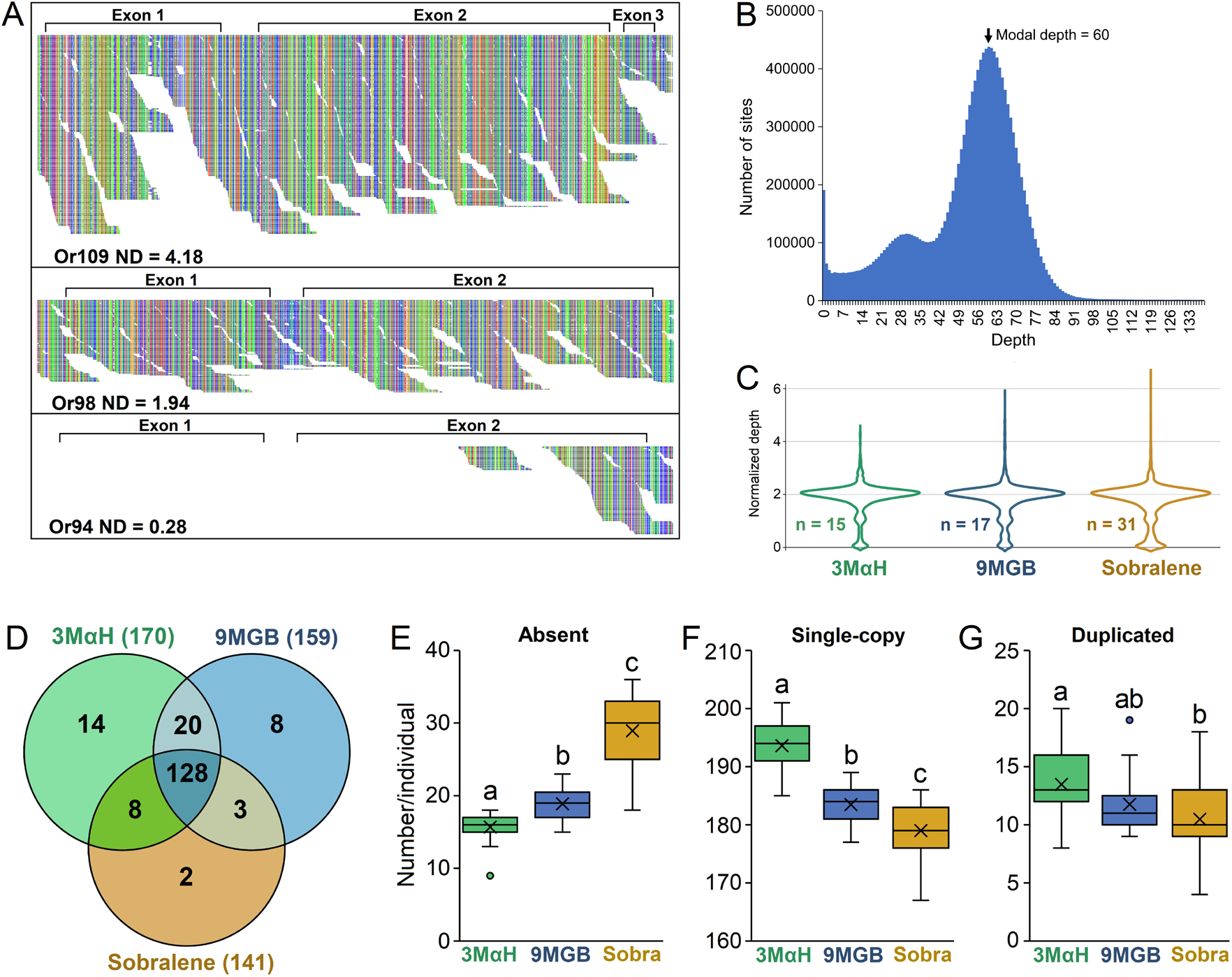
Visualization of reads aligned to chemoreceptor loci revealed variation in coverage that indicated potential copy number variation. **(A)** For example, JAC01 Or109 had much deeper coverage than Or98, while Or94 had reads mapped only at the end of the second exon. The Tablet software program was used for visualization of the mapped reads. **(B)** To quantify these differences for comprehensive analysis of all 245 chemoreceptor loci, we calculated background-normalized sequencing depth of each gene using the modal depth across the exons in all protein coding genes. **(C)** A central tendency of ~2 is expected for single copy genes with two intact alleles. Normalized depth was rounded to the nearest whole number as a proxy for copy number (CN). (D) The number of intact chemoreceptors (CN≥2) in all individuals of a chemotype ranged from 141 (Sobralene) to 170 (3MαH). **(E)** The mean number of absent (CN = 0) genes differed among all chemotypes (*P* <0.001), with Sobralene individuals having the most and 3MαH individuals having the fewest. **(F)** Accordingly, the number of single-copy (CN = 2) genes differed among all three chemotypes (*P* <0.001), with 3MαH individuals having the largest number and Sobralene individuals having the fewest. **(G)** The number of duplicated genes (CN>2) differed only 3MαH and sobralene individuals.

To validate our method for estimating gene copy number, we annotated a gene model or confirmed absence for 522 chemoreceptor genes in the *de novo* assemblies (Supplemental Table S5). We were unable to annotate any intact genes when CN=0. Annotation of the duplications was not always possible, most likely due to high sequence similarity causing them to be collapsed during *de novo* assembly. Within-population CNV was apparent at several loci, evidenced by CN of approximately 2, 1, and 0, which we inferred as two intact alleles (A^+/+^), one intact and one degraded allele (A^+/-^), and two degraded alleles (A^-/-^), respectively, which is further supported by sites with heterozygosity in A^+/+^, but not in A^+/-^ (Fig S5A). However, not all genes with CN=1 represented a combination of an intact and a degraded allele. Some genes were degraded in regions of a gene, with approximately half of the gene intact (Fig S5B).

Of the 245 genes in the reference chemoreceptor genome, only 100 were single-copy (CN=2) in all 63 individuals, while those that were at least single-copy (CN≥2) in each of the three chemotypes ranged from 141 in Sobralene to 170 in 3MαH, with only 128 shared among all individuals (Fig 3D). Further, the mean number of absent (CN=0) genes differed among the three chemotypes (p<0.001) with Sobralene individuals having 28.9 ± 4.5 (mean ± SD), followed by 9MGB with 18.9 ± 2.2, and 3MαH with 15.7± 2.3 (Fig 3E). Accordingly, the mean number of single-copy genes (CN=2) differed among chemotypes (p<0.001), with 3MαH (193.6 ±4.3) followed by 9MGB (183.5 ±3.5) and Sobralene (179 ± 4.6). The mean number of duplicated genes (CN>2) was greatest in individuals from 3MαH (13.5 ± 3.2) compared to 9MGB (11.8 ± 2.7) and Sobralene (10.5 ± 3.0) which did not significantly differ (Fig 3 E-G).

Hierarchical analysis and PCA of CN grouped individuals according to their putative chemotype (Fig 4A and B). PC1 and PC2 explained 27.4% and 17.3% of the variation, respectively (Fig 4B). Of the three chemoreceptor gene families, the ORs were the most dynamic with 72.1% displaying CNV, followed by 46.3% of GRs and 21.7% of IRs.

**Fig. 4.**
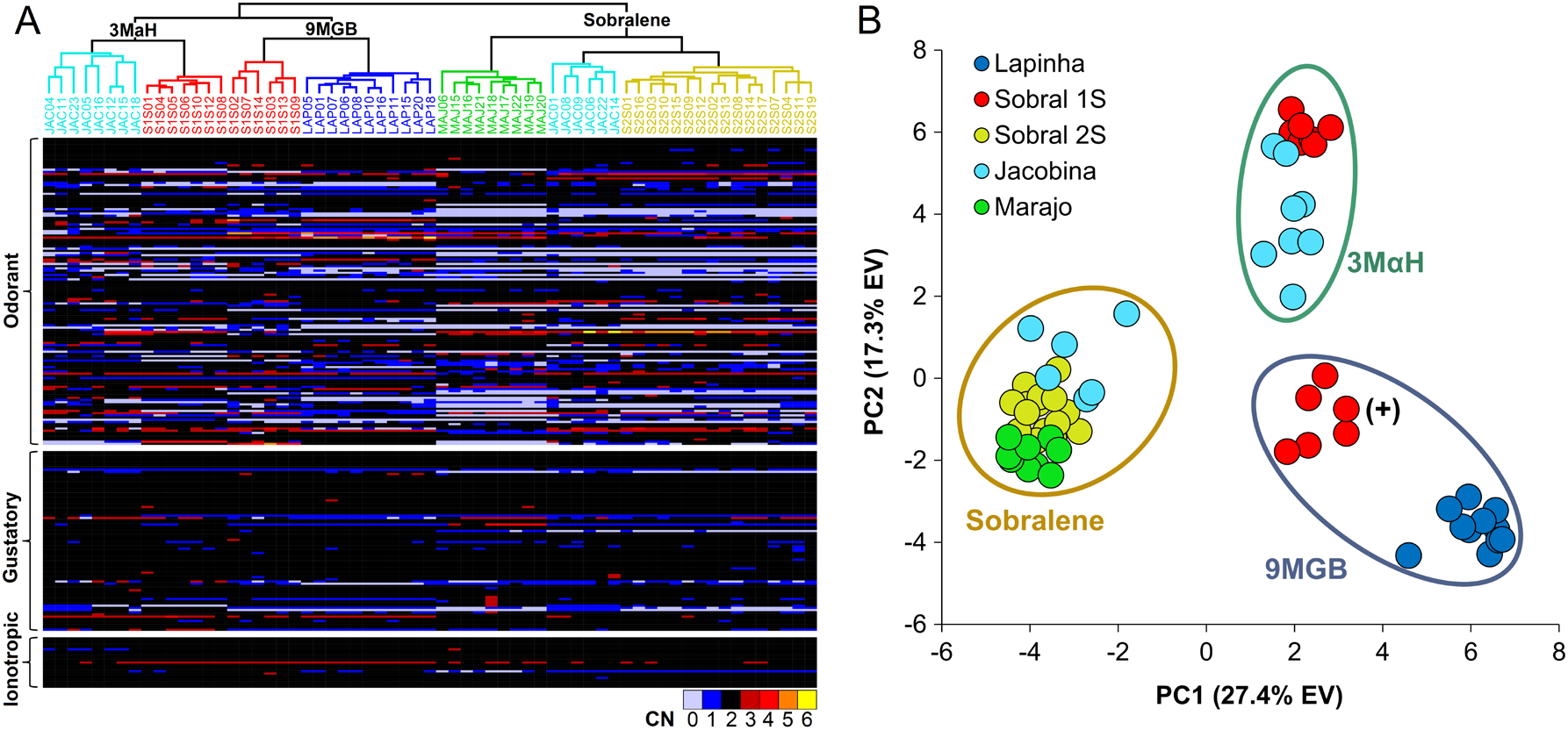
Relationships among 63 *L. longipalpis* from four sites in Brazil (Lapinha Cave, Sobral, Jacobina and Marajó) based on copy number (CN) of 245 chemoreceptor genes. **(A)** Hierarchical analysis of gene CN clustered individuals according to their putative chemotype as determined previously using SNPs. A heatmap of CN illustrates the large number of genes with CNV among the odorant receptors. **(B)** PCA of gene CN showed a similar pattern, wherein Sobral 2S (2 spots), Marajó and six Jacobina clustered together (Sobralene); seven Sobral 1S (1 spot) and eight Jacobina clustered together (3MαH); and Lapinha Cave and six Sobral 1S (one spot) were in separate clusters (9MGB), which is consistent with the genetic differentiation observed by Hamilton et al. (2005) thus leading them to classify these as different chemotypes (9MGB and 9MGB+) due to the larger quantity of 9MGB produced by males from Lapinha [10].

We used pairwise V_ST_ to identify genes exhibiting the greatest CNV between chemotypes. Of the 60 genes with V_ST_ >0.5 in all three pairwise analyses, 43 are ORs, 16 are GRs, and only one is an IR (Fig 5 A–C). In most cases, genes with V_ST_ >0.5 had CN<2 in some individuals from both chemotypes. That is, few were intact (CN≥2) in all individuals of a chemotype, suggesting that they were not under strong purifying selection in either chemotype. However, several genes were intact in one chemotype and largely absent in the other. In the pairwise analysis of 9MGB and Sobralene, Or37, Or43, Or115 and Or139 have CN≥2 in all 9MGB, but CN<2 in some of the Sobralene individuals. In the pairwise analysis between 3MαH and Sobralene, Or115 and Gr37 have a CN≥2 in all 3MαH, but are absent in some Sobralene (Fig 5B). In the analysis between 9MGB and 3MαH, Or23, Or116 and Or118 are intact only in 9MGB, while Or14, Or43 and Gr71 are intact only in 3MαH (Fig 5C). Or115 and Or116 are noteworthy because they are paralogs and were among the highest V_ST_ in the pairwise analyses (Fig 5 A–C). Furthermore, they are in a neighboring clade to the *Mayetiola destructor* and *D. melanogaster* pheromone receptors (Fig S1). While annotating Or115 and Or116 in the *de novo* assemblies, a third paralog (Or137) was found in all of the individuals, but was missing in the reference assembly (Supplemental Table S5).

**Fig. 5.**
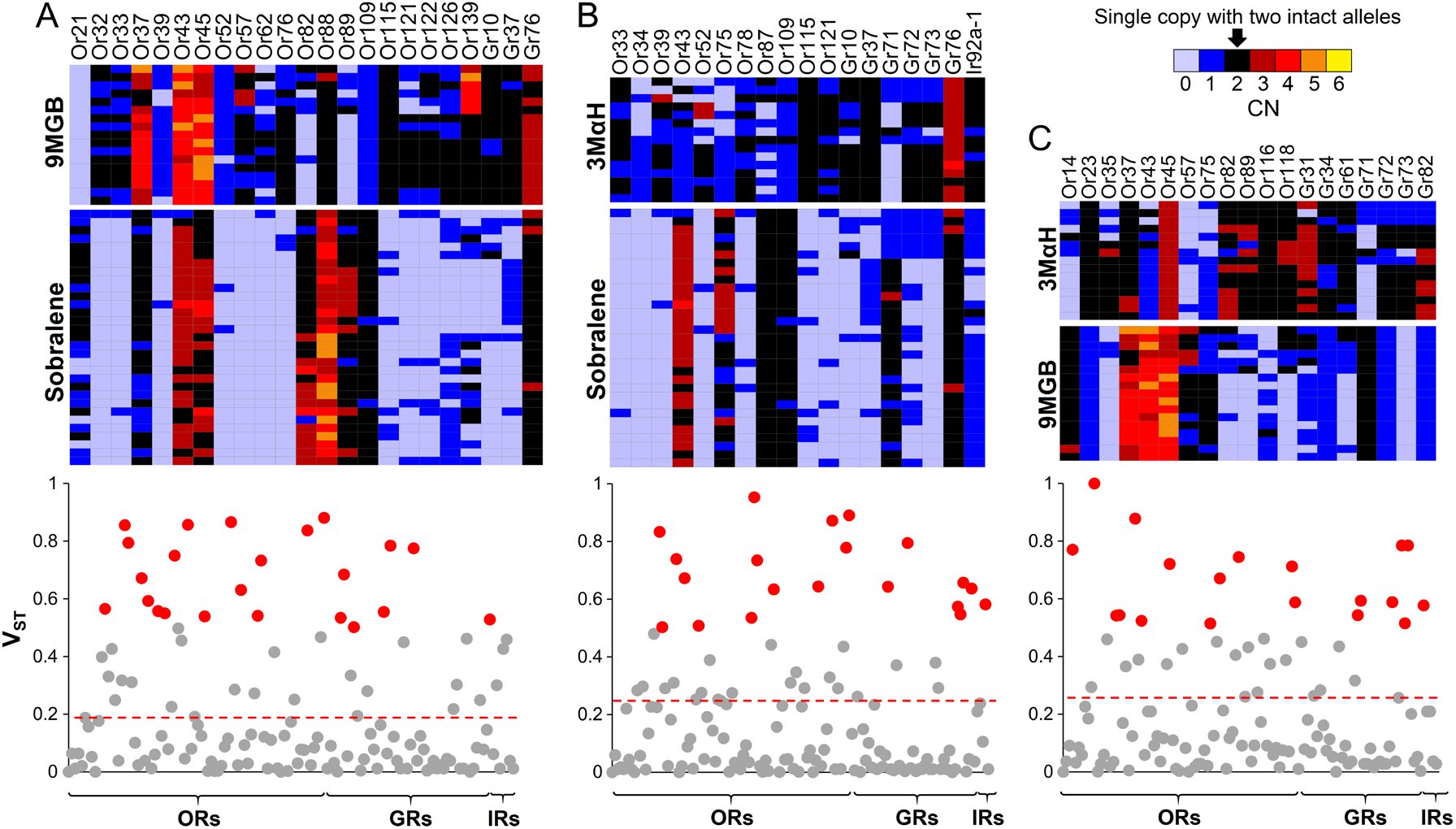
V_ST_ was used to identify the most differentiated genes based on CNV. Heatmaps of copy number (CN) and Manhattan plots of V_ST_ between **(A)** sobralene and 9MGB, **(B)** sobralene and 3MαH, and **(C)** 9MGB and 3MαH (V_ST_ >0.5 highlighted in red). The dashed lines indicate the threshold for significance (0.99) based on 1,000 permutations. Heatmaps illustrate CN of genes with V_ST_>0.5. Of the 60 genes with V_ST_>0.5 in all three pairwise analyses, 43 were ORs, 16 were GRs and only one was an IR.

## Discussion

Evidence of the genetic variability and its potential implication in vector management strategies was established early in sand flies: biting of *L. longipalpis* females from Costa Rica did not leave long-lasting erythemas that are characteristic in Brazil and Colombia even though parasites were indistinguishable from those locations [8, 54]. In addition, the population structure and genetic variability within and among different sand fly populations was found to influence vectorial capacity [55]. Consequently, an array of molecular and biochemical markers have been explored to identify the genotypes underlying phenotypes of interest, such as vector competence, that should be considered during the planning and implementation of integrated control strategies for leishmaniasis [56, 57].

Since the initial interactions between an organism and its chemical landscape are mediated by chemoreceptors, the role of these proteins in ecological adaptation is paramount. Genomic divergence in the form of SNPs and CNV is pervasive in chemosensory gene families in both mammals and insects [27, 58, 59]. Birth-and-death evolution of these gene families often manifests as divergent, taxonomically restricted gene lineages, which have been implicated in the evolution of sociality in the honey bee [60]. The sand flies present an extreme example among the Diptera, with over 80% of their ORs belonging to a single, highly expanded lineage. Aside from a few exceptions, the *P. papatasi* and *L. longipalpis* reference chemoreceptor genomes are similar in size and content (Fig 1A), therefore, the amount of CNV among and within chemotypes in Brazil was unexpected. Our results indicate differential CNV among the chemoreceptors wherein ORs were the most dynamic, followed by the GRs and IRs (Fig 3). Similar observations are noted in pea aphids that were attributed to strong drift and selection [61]. A major role for CNV in adaptative innovation was proposed nearly 50 years ago [53]. Unequal crossover during meiosis is thought to be the primary mechanism leading to CNV in the insect chemoreceptors [25, 27]. One of the outcomes of these duplications is the process of neofunctionalization, the development of novel function due to relaxed constraints on a paralog [62]. The adaptive radiation model of neofunctionalization predicts that the emergence of a duplicated gene produces additional variation through relaxed selection, which is followed by “competitive evolution” where the most favorable variant is preserved. Sexual communication in sand flies herein offers an exciting model to test these hypotheses.

Sex pheromones mediate intraspecific communication in many systems [63], and mediate behavioral isolation between insect populations [64]. In sand flies, two different chemotypes in São Paulo (9MGB and sobralene) showed that the expansion of the visceral leishmaniasis disease (canine and human form) was only correlated with the dispersion route of the 9MGB chemotype [65]. Further analyses of the variation in chemical profiles of *Lutzomyia* populations [6] are warranted, and correlating this with both geographic location and genomic composition will enhance our understanding of the evolution of chemical communication. Pheromone detecting ORs among the Diptera are not well known. In the fruit fly, DmelOr67d detects the male pheromone cVA [66], which is phylogenetically similar to the hessian fly pheromone receptor *Mdes115* [67] (Fig 1S). Our analysis identifies a few promising candidate ORs from the sand flies that cluster with these pheromone receptors (Fig 1S). Identification and functional characterization of the aggregation-sex pheromone receptors in *L. longipalpis* will provide insights into the evolutionary mechanisms associated with assortative mating between chemotypes.

In addition to aggregation-sex pheromones, the molecular basis of other sand fly behaviors, such as host seeking, oviposition and sugar feeding, could provide insights into these life history traits. Several studied have investigated host odors in combination with aggregation-sex pheromones and have suggested additive and, or synergistic effects [6, 68]. Members of the *L. longipalpis* species complex have an irregular distribution and have adapted to a variety of tropical habitats, ranging from rocky and arid to humid and forested areas. Consequently, insights from their chemoreceptor genome, especially IRs that are being increasingly implicated in diverse roles in environmental sensing [69], will be essential to understanding the role of chemosensory gene family evolution in sexual communication and ecological adaptation.

Historically, some of the most successful campaigns against vector-borne diseases have been those targeted against the vectors [70]. Control of leishmaniasis has largely depended on reducing human exposure to sand flies using residual insecticides, repellents, bed nets and population control strategies [71]. However, there is a growing interest in developing novel vector management strategies by exploiting the chemosensory behaviors of the vector insects [72]. Our data provides novel insights into complex population structure in Brazilian sand flies, and highlights the role of chemoreceptors in the evolution of novel pheromone types, thus adding to the theoretical framework of speciation by sexual selection.

## Author Contributions

P.V.H., M.A.M. and Z.S. designed research; P.V.H., N.T., C.N.S. and Z.S. performed research; P.V.H., N.T., RJ.N., F.L., Y.L., A.D.N., M.A.M., C.N.S. and Z.S. participated in analyzing data; P.V.H. and Z.S. wrote the paper.

## Acknowledgements

We thank Hugh M. Robertson (University of Illinois, Urbana-Champaign, IL) for his guidance on annotating the chemoreceptors, and David C. Rinker (Vanderbilt University, TN) for his recommendations for calculating gene copy number and V_ST_. We would also like to thank Cleilton Sampaio de Farias (Instituto Federal do Acre, Brazil) for the map construction, and Felipe Vigoder (Universidade Federal do Rio de Janeiro, Brazil) for sharing the original recordings of the copulation songs displayed in Fig 2. We acknowledge the late Alexandre Peixoto, whose previous work inspired this research and who participated in the sampling strategy prior to his untimely death. This research work was supported by funding from National Institute of Food and Agriculture, US Department of Agriculture (under HATCH Project 2353077000). *L. longipalpis* genome data were generated by Baylor College of Medicine Human Genome Sequencing Center in collaboration with Washington University in Saint Louis as part of the sand fly NHGRI project. Genome data can be obtained at www.vectorbase.org.

