## Supplemental figures for "Molecular signatures of sexual communication in the phlebotomine sand flies"

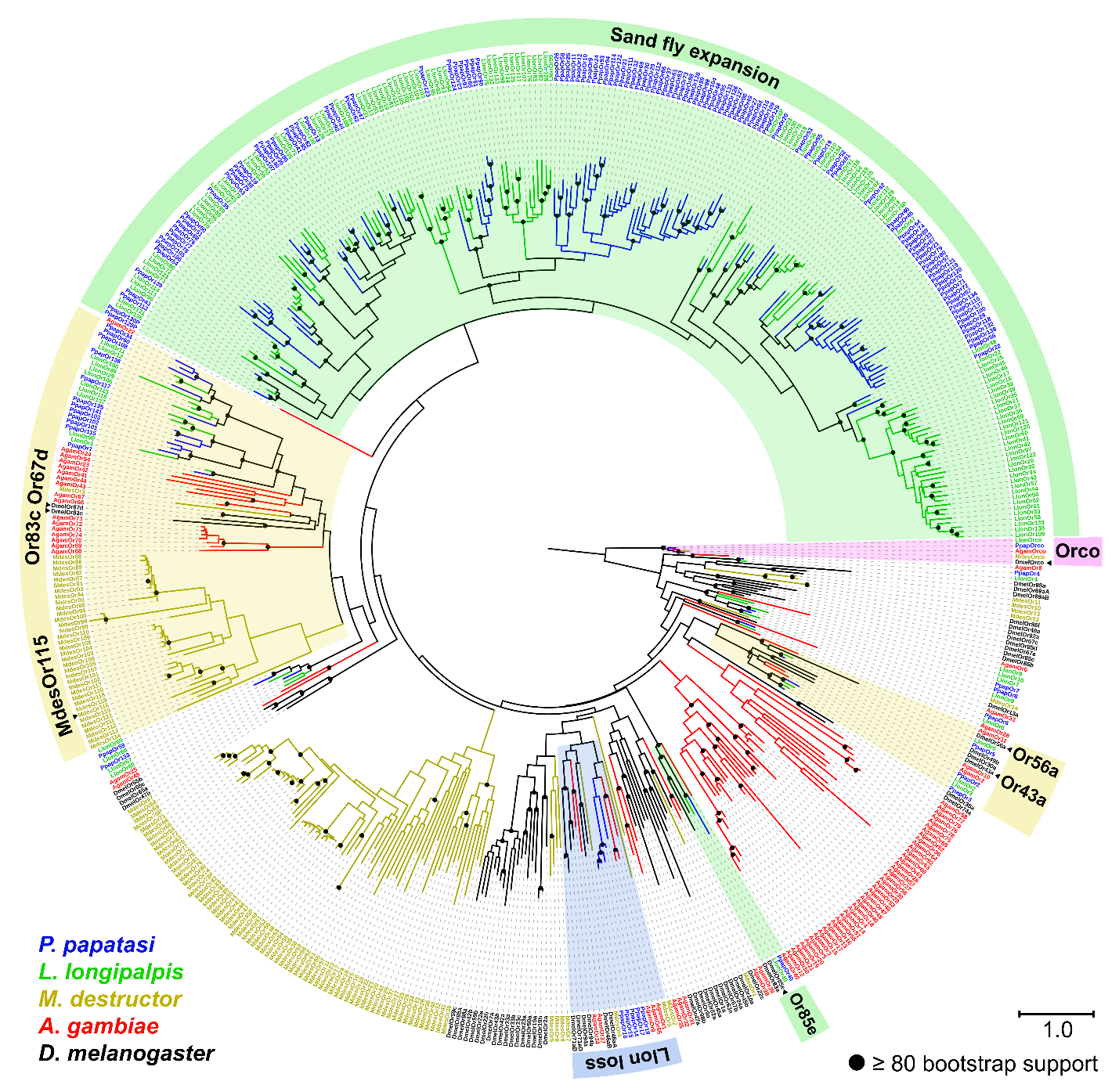

**Fig. S1.** Phylogenetic relationships among ORs in *L. longipalpis, P. papatasi, A. gambiae, M. destructor* and *D. melanogaster*. Expanded, conserved and lost OR lineages are shaded. The phylogeny was estimated using the JTT model of protein substitution and Maximum Likelihood method in RAxML v.8.2.4 [1] . The tree is rooted at the branch leading to Orco. Branch support based on 500 bootstrap replications.

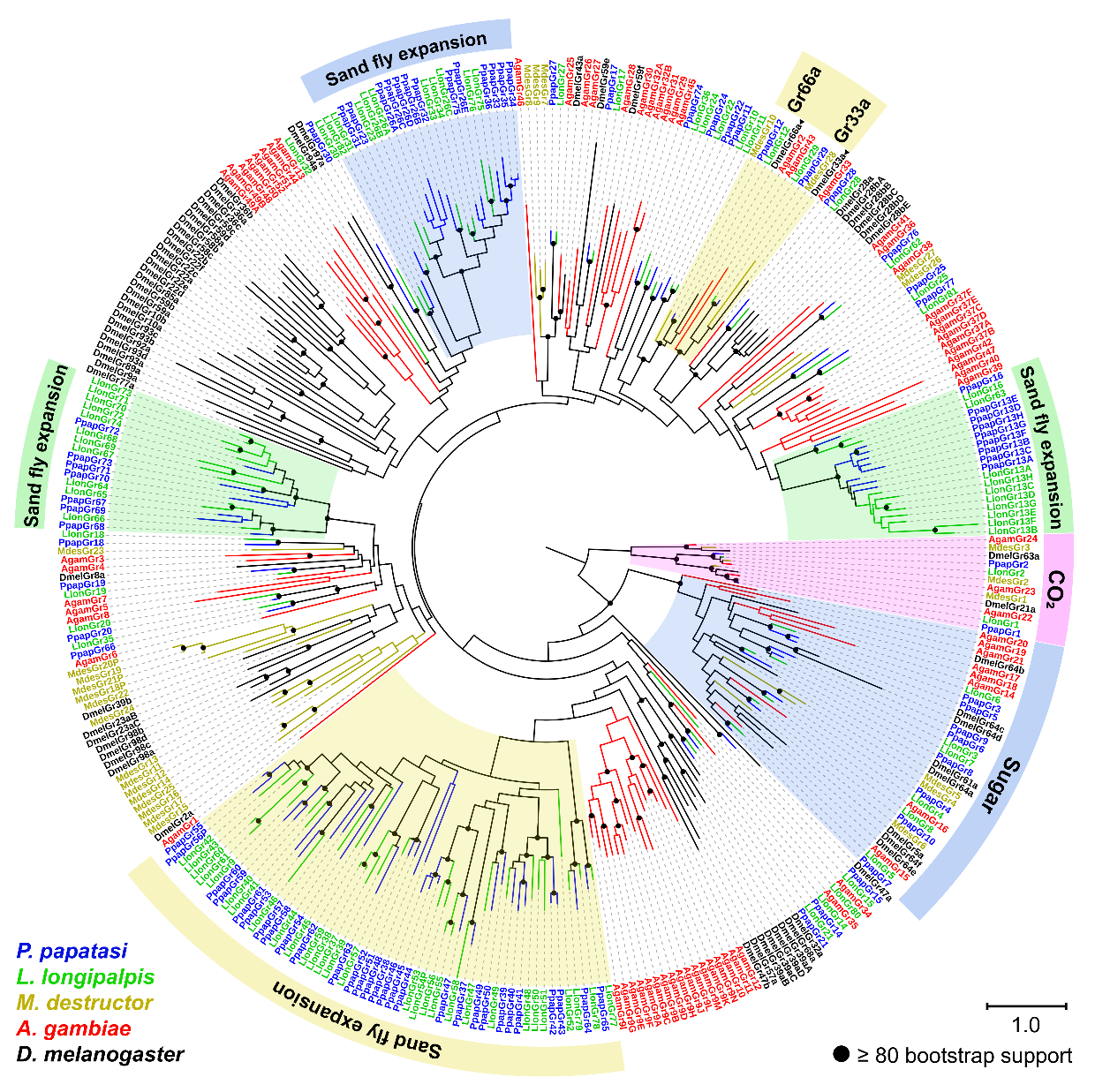

**Fig. S2.** Phylogenetic relationships among GRs in *L. longipalpis, P. papatasi, A. gambiae, M. destructor* and *D. melanogaster*. Expanded and conserved GR lineages are shaded. The phylogeny was estimated using the JTT model of protein substitution and Maximum Likelihood method in RAxML v.8.2.4 [1]. The tree is rooted at the branch leading to the CO_2_ receptors. Branch support based on 500 bootstrap replications.

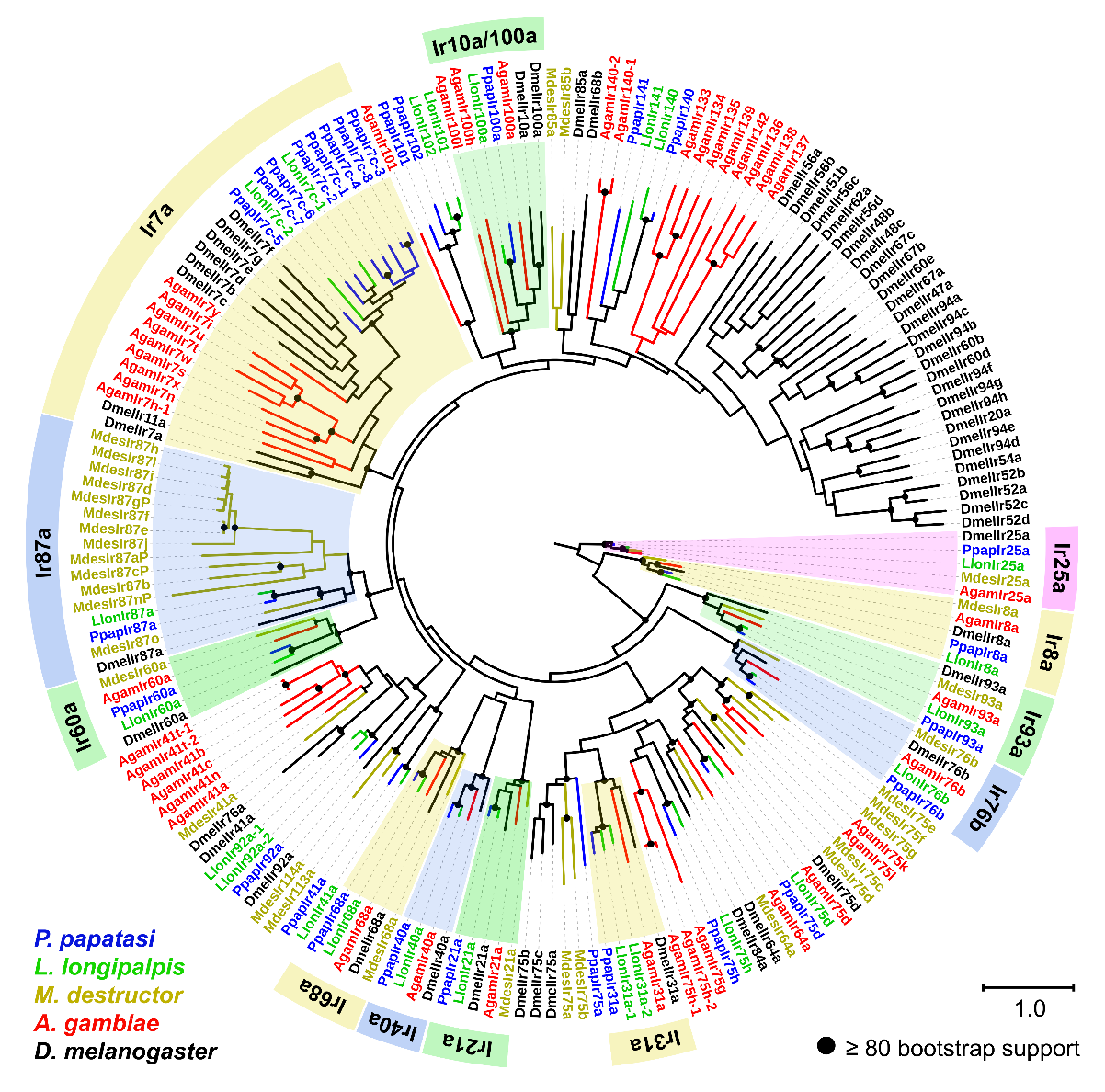

**Fig. S3.** Phylogenetic relationships among IRs in in *L. longipalpis,* *P. papatasi*, *A. gambiae,* *M. destructor* and *D. melanogaster.* Conserved IR lineages are shaded. The phylogeny was estimated using the JTT model of protein substitution and Maximum Likelihood method in RAxML v.8.2.4 [1]. The tree is rooted at the branch leading to Ir25a and Ir8a. Branch support based on 500 bootstrap replications.

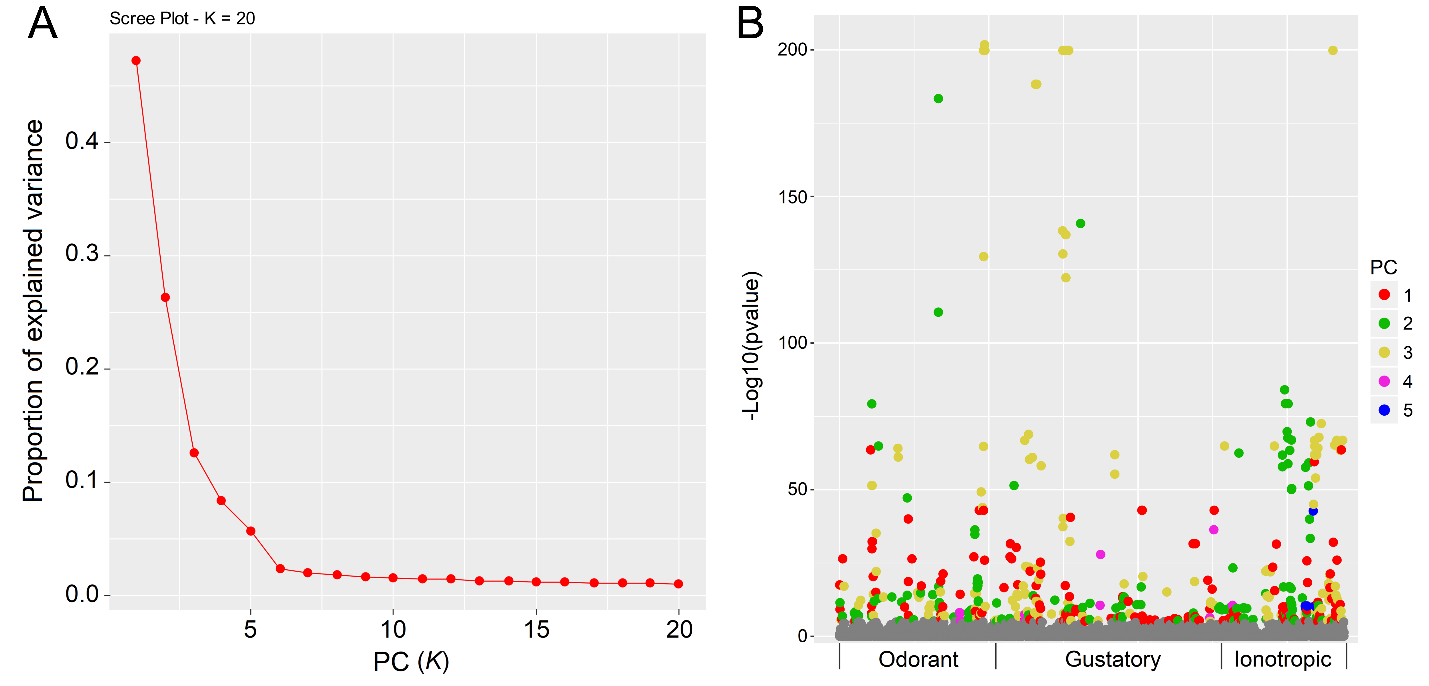

**Fig. S4.** *pcadapt* was used to identify loci associated with population structure based on SNPs in the exons of 245 chemoreceptor genes in 63 individuals. (A) An optimal number of PCs, *K* = 5, was selected following preliminary analysis with *K* = 20 and subsequent visualization of the scree plot. (B) Scatterplot showing the outliers after Bonferroni correction (0.05) and the PCs they are associated with based on component-wise analysis in PCadapt [2].

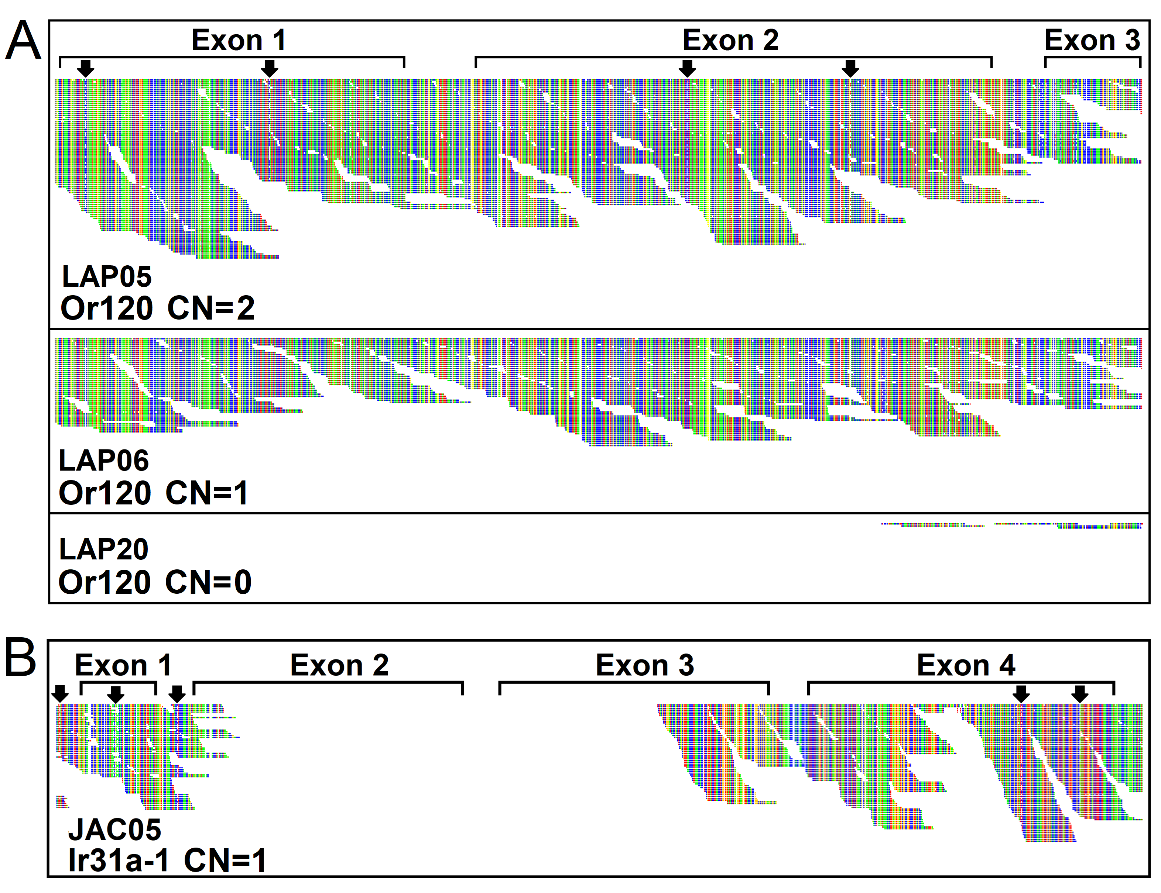

**Fig. S5.** Two distinct conditions were identified producing a copy number of 1 (CN=1), which is half the expected CN for a single-copy gene with two intact alleles. **(A)** Or120 has CN=2 in LAP05, CN=1 in LAP06 and CN=0 in LAP20, suggesting the presence of intrapopulation variation in the form of two intact alleles (A^+/+^), an intact and a degraded allele (A^+/-^), and two degraded alleles (A^-/-^), respectively. Arrows indicate sites of heterozygosity in LAP05, while all sites are homozygous in LAP06. **(B)** Ir31a-1 in JAC05 illustrates the second situation where CN=1 represents a moderately degraded pseudogene. Arrows indicate sites of heterozygosity indicating the parents likely had at least one allele with a similar degree of degradation. The Tablet software program [3] was used for visualization of read alignments.
