## Supplemental Table legends for "Molecular signatures of sexual communication in the phlebotomine sand flies"

**Supplemental Table 1S.** Accession numbers for SRAs and assigned chemotypes for the 63 *L. longipalpis* individual genome sequences used in the study.

**Supplemental Table 2S.** Chemoreceptor gene models for ORs, GRs and IRs for *L. longipalpis* and *P. papatasi*.

**Supplemental Table 3S.** SNPs in the 100 single-copy orthologs among the *L. longipalpis* field collections.

**Supplemental Table 4S.** Predicted gene copy number for 245 chemoreceptor genes in 63 *L. longipalpis* individuals.

**Supplemental Table 5S.** Gene models and confirmed absences for 522 genes based on manual annotations of the 63 *de novo* assemblies.
