## Supplemental Table S1 for "Molecular signatures of sexual communication in the phlebotomine sand flies"

**Table S1.** SRA identification numbers, collection sites and putative chemotypes for *L. longipalpis* individuals used in the study. Sobral 1-spot and 2-spot are designated S1S and S2S, respectively.

| SRA | Individual | Group | Collection site | Chemotype |
| --- | --- | --- | --- | --- |
| SRX1537447 | S1S07 | S1S-A | Sobral | 9MGB+ |
| SRX1537448 | S1S09 | S1S-A | Sobral | 9MGB+ |
| SRX1537451 | S1S14 | S1S-A | Sobral | 9MGB+ |
| SRX1537452 | S1S13 | S1S-A | Sobral | 9MGB+ |
| SRX1537454 | S1S03 | S1S-A | Sobral | 9MGB+ |
| SRX1537455 | S1S02 | S1S-A | Sobral | 9MGB+ |
| SRX1537442 | S1S06 | S1S-B | Sobral | 3MαH |
| SRX1537444 | S1S05 | S1S-B | Sobral | 3MαH |
| SRX1537445 | S1S12 | S1S-B | Sobral | 3MαH |
| SRX1537446 | S1S10 | S1S-B | Sobral | 3MαH |
| SRX1537449 | S1S04 | S1S-B | Sobral | 3MαH |
| SRX1537453 | S1S08 | S1S-B | Sobral | 3MαH |
| SRX1537456 | S1S01 | S1S-B | Sobral | 3MαH |
| SRX1537417 | LAP08 | LAP | Lapinha | 9MGB |
| SRX1537418 | LAP16 | LAP | Lapinha | 9MGB |
| SRX1537419 | LAP05 | LAP | Lapinha | 9MGB |
| SRX1537420 | LAP07 | LAP | Lapinha | 9MGB |
| SRX1537423 | LAP20 | LAP | Lapinha | 9MGB |
| SRX1537424 | LAP10 | LAP | Lapinha | 9MGB |
| SRX1537427 | LAP06 | LAP | Lapinha | 9MGB |
| SRX1537428 | LAP11 | LAP | Lapinha | 9MGB |
| SRX1537429 | LAP18 | LAP | Lapinha | 9MGB |
| SRX1537430 | LAP01 | LAP | Lapinha | 9MGB |
| SRX1537431 | LAP15 | LAP | Lapinha | 9MGB |
| SRX1537432 | MAR19 | MAR | Marajo | Sobralene |
| SRX1537434 | MAR17 | MAR | Marajo | Sobralene |
| SRX1537435 | MAR16 | MAR | Marajo | Sobralene |
| SRX1537436 | MAR21 | MAR | Marajo | Sobralene |
| SRX1537437 | MAR15 | MAR | Marajo | Sobralene |
| SRX1537438 | MAR20 | MAR | Marajo | Sobralene |
| SRX1537439 | MAR22 | MAR | Marajo | Sobralene |
| SRX1537440 | MAR06 | MAR | Marajo | Sobralene |
| SRX1537441 | MAR18 | MAR | Marajo | Sobralene |
| SRX1537458 | S2S07 | S2S | Sobral | Sobralene |
| SRX1537459 | S2S02 | S2S | Sobral | Sobralene |
| SRX1537460 | S2S04 | S2S | Sobral | Sobralene |
| SRX1537461 | S2S08 | S2S | Sobral | Sobralene |
| SRX1537462 | S2S03 | S2S | Sobral | Sobralene |
| SRX1537463 | S2S15 | S2S | Sobral | Sobralene |
| SRX1537464 | S2S14 | S2S | Sobral | Sobralene |
| SRX1537465 | S2S16 | S2S | Sobral | Sobralene |
| SRX1537466 | S2S19 | S2S | Sobral | Sobralene |
| SRX1537467 | S2S13 | S2S | Sobral | Sobralene |
| SRX1537468 | S2S09 | S2S | Sobral | Sobralene |
| SRX1537469 | S2S17 | S2S | Sobral | Sobralene |
| SRX1537470 | S2S01 | S2S | Sobral | Sobralene |
| SRX1537471 | S2S11 | S2S | Sobral | Sobralene |
| SRX1537472 | S2S10 | S2S | Sobral | Sobralene |
| SRX1537473 | S2S12 | S2S | Sobral | Sobralene |
| SRX1537403 | JAC23 | JAC-A | Jacobina | 3MαH |
| SRX1537405 | JAC05 | JAC-A | Jacobina | 3MαH |
| SRX1537407 | JAC11 | JAC-A | Jacobina | 3MαH |
| SRX1537409 | JAC12 | JAC-A | Jacobina | 3MαH |
| SRX1537410 | JAC04 | JAC-A | Jacobina | 3MαH |
| SRX1537411 | JAC15 | JAC-A | Jacobina | 3MαH |
| SRX1537413 | JAC16 | JAC-A | Jacobina | 3MαH |
| SRX1537415 | JAC18 | JAC-A | Jacobina | 3MαH |
| SRX1537404 | JAC08 | JAC-B | Jacobina | Sobralene |
| SRX1537406 | JAC22 | JAC-B | Jacobina | Sobralene |
| SRX1537408 | JAC09 | JAC-B | Jacobina | Sobralene |
| SRX1537412 | JAC01 | JAC-B | Jacobina | Sobralene |
| SRX1537414 | JAC06 | JAC-B | Jacobina | Sobralene |
| SRX1537416 | JAC14 | JAC-B | Jacobina | Sobralene |
